# The neuronal associations of respiratory-volume variability in the resting state

**DOI:** 10.1101/2020.10.01.322800

**Authors:** Sayedmohammad Shams, Pierre LeVan, J. Jean Chen

## Abstract

The desire to enhance the sensitivity and specificity of resting-state (rs-fMRI) measures has prompted substantial recent research into removing noise components. Chief among contributions to noise in rs-fMRI are physiological processes, and the neuronal implications of respiratory-volume variability (RVT), a main rs-fMRI-relevant physiological process, is incompletely understood. The potential implications of RVT in modulating and being modulated by autonomic nervous regulation, has yet to be fully understood by the rs-fMRI community. In this work, we use high-density electroencephalography (EEG) along with simultaneously acquired RVT recordings to help address this question. We hypothesize that (1) there is a significant relationship between EEG and RVT in multiple EEG bands, and (2) that this relationship varies by brain region. Our results confirm our first hypothesis, although all brain regions are shown to be equally implicated in RVT-related EEG-signal fluctuations. The lag between RVT and EEG is consistent with previously reported values. However, an interesting finding is related to the polarity of the correlation between RVT and EEG. Our results reveal potentially two main regimes of EEG-RVT association, one in which EEG leads RVT with a positive association between the two, and one in which RVT leads EEG but with a positive association between the two. We propose that these two patterns can be interpreted differently in terms of the involvement of higher cognition. These results further suggest that treating RVT simply as noise is likely a questionable practice, and that more work is needed to avoid discarding cognitively relevant information when performing physiological correction rs-fMRI.

## Introduction

The involvement of physiological processes in the resting-state fMRI signal and in the resulting functional connectivity measures is an important question, and has yet to be fully understood. Physiological bias in resting-state fMRI is a debated topic (Liu et al., 2017; Yan et al., 2009). Thus far, the most substantial contributors to the resting-state fMRI signal relevant to functional connectivity measurements are thought to be respiratory volume per unit time (RVT) variability, heart-rate variability (HRV) and end-tidal CO_2_ (PETCO_2_) fluctuations.

The relationship between HRV and resting-state brain activity as well as functional connectivity is well studied (Ako et al., 2003; Alba et al., 2019; Chang et al., 2016; Fan et al., 2012; Jennings et al., 2016; Kuo et al., 2016; Sakaki et al., 2016). The relationship between PETCO_2_ fluctuations and neural activity has been investigated using both electroencephalography (EEG) and magnetoencephalography (MEG) (Driver et al., 2016; Xu et al., 2011). However, the relationship between RVT and neural activity remains unclear.

RVT is one of the main physiological fluctuations that is regarded as noise in rs-fMRI, but this notion has been questioned (Iacovella and Hasson, 2011). Indeed, the respiratory cycle has beens related to arousal as measured using EEG (Busek and Kemlink, 2005)(Morelli et al., 2018). RVT is also a marker of the autonomic nervous system (Chung et al., 2019). However, in the context of resting-state fMRI, so far only one study, by Yuan et al., addresses this question directly. Yuan et al. observed that the alpha-EEG-RVT association is stronger during eyes-closed resting state (Yuan et al., 2013). Furthermore, alpha global-field potential (GFP) and respiration fluctuations were found to be correlated in several regions in a highly consistent spatial pattern, including the visual/parietal cortex, superior/middle temporal gyrus, inferior frontal gyrus, inferior parietal lobule, thalamus and caudate. Notably, the previous work targeted only the alpha band (frequency between 7.5 and 13 Hz), which appears when closing the eyes and relaxing, and disappears when opening the eyes or alerting by any mechanism (thinking, calculating). Beyond the alpha band, the following frequency bands are also implicated in cognition and arousal:

- Delta: typically the dominant rhythm in stages 3 and 4 of sleep. It is usually most prominent frontally in adults.
- Theta: classified as “slow” activity. It is typically not prominent in awake adults.
- Beta: classified as fast activity, usually seen on both sides in symmetrical distribution and is most evident frontally. It is accentuated in the eyes-open state.

These frequency bands also have differential spatial distributions. For instance, the alpha band displays the highest power in the occipital lobe. Moreover, there has been no replication study since the work of Yuan et al., and no follow up that extends the investigation. Thus, in this study, we aim to expand the knowledge of the neural associations of RVT in two directions: (1) inclusion of the delta, theta and beta bands in addition to alpha; (2) distinction amongst different cortical regions. We hypothesize that: (1) significant relationships exist between RVT and EEG, spanning multiple frequency bands; (2) these relationships vary across different brain regions.

## Methods

### EEG acquisition

The EEG data were acquired from seventeen healthy volunteers using a 256-channel MR-compatible EEG system with sponge-based electrodes (EGI; sampling rate 1000 Hz). The data were hardware filtered to be between 0.1-400 Hz. Subjects were instructed to keep their eyes closed and be vigilant. The recordings lasted 5 min each and were performed inside a MRI environment (Siemens TIM Trio 3 Tesla) as part of a separate EEG-fMRI study. During the recordings, a MR-compatible camera (Metria Innovation Inc., Milwaukee, USA), was fixed inside the MRI scanner bore to track the position of a marker (Maclaren et al., 2012) fixed to the base of the subject’s forehead. The camera software then yielded full position and orientation information with 6 degrees of freedom and a sampling rate of 80 Hz, which was used for EEG preprocessing, as described later.

### Physiological Data

During each recording session, ECG was also recorded using the MRI-scanner’s built-in electrodes (Siemens, Erlangen, Germany). A pneumatic belt positioned at the level of the abdomen was used to measure the respiration during the data collection. In this study we exploited the respiration volume per time (RVT) as the characterization of the respiratory fluctuations according to Birn et al. (2006). To assess respiration volume per time (RVT), we used the open source TAPAS physio toolbox available at https://www.tnu.ethz.ch/en/software/tapas.html (Kasper et al., 2017).

### EEG data preprocessing

Then, preprocessing was performed to remove gradient and ballistocardiographic (BCG) artefacts due to the MRI environment, taking advantage of the camera-recorded motion parameters to achieve an optimal denoising performance. Specifically, these parameters were used to model modulations of the gradient artifact across time, as described in (LeVan et al., 2016). As for the BCG artifact, it was modelled as a filtered version of the measured head position changes by regressing out successively lagged copies of the motion parameters (LeVan et al., 2013). This was followed by commonly used average-artifact subtraction (AAS) to remove residual artifacts. Furthermore, we applied independent component analysis (ICA) on the EEG time series and manually identified the independent components (ICs) that are related to the residual BCG artifacts and movement effects to discard them from the analysis (Luo et al., 2010).

We have also added an automated preprocessing step to specify the highly contaminated channels for each recording and discard them from the analysis. To do so, we used a short-time Fourier Transform (1-sec sliding Hanning window with step = 100 ms) and generated the power spectrogram for each channel separately. For each channel, the power spectrum of the frequencies greater than 15 Hz was compared to the power spectrum of the frequencies less than 14 Hz. We empirically observed that noisy channels, notably due to residual gradient artifacts or poor electrode connections, had high-frequency power exceeding around 20% of the low frequency power. Therefore, all channels where the high-frequency power above this threshold were discarded from the analysis.

### Estimation of EEG-Bands Fluctuations

The previously performed time-frequency analysis was used to estimate the power fluctuation of different EEG frequency bands. The power spectrum for each electrode time series was calculated in intervals of 100 ms. At each time point, the EEG-bands power were obtained from the spectrogram by averaging over the frequency ranges corresponding to each EEG band: 0.5 −3 Hz for delta, 3.5-7.5 Hz for theta, 8-12 Hz for alpha, and 13-30 Hz for beta. As the signal-to-noise ratio (SNR) in the gamma band was not reliable in this resting-state design, we excluded the gamma band from analysis.

For each frequency band, we identified subject-specific peaks (maxima) and associated peak frequencies. For instance, for the alpha band, to accurately isolate the frequency range for alpha band, we first extracted the peak frequency for alpha band (maximum power in 8-12 Hz) and for each subject, and then used a bandwidth of 2 Hz around the extracted band peak frequency for each subject separately (similar to what was done in (Yuan et al., 2013)). To obtain each band’s global field power (GFP) and lobe specific field power (LSFP), we applied principal component analysis (PCA) over the power spectrum fluctuation time series of all channels or channels corresponding to each lobes and used the first component of the PCA as the GFP and LSFP, respectively. In order to specify the electrodes that correspond to each lobe, we first pre-determined the representative electrodes for each lobes as follows:

- Occipital: O1, Oz, O2
- Parietal: P3, Pz, P4
- Frontal: F7, F3, Fz, F4, F8

The topographic maps of the distribution of the spatial locations of the electrodes assigned to frontal, parietal, and occipital lobes are shown in Fig. 1, in which the spatial distributions resulted from the algorithm based on the distance of each electrode and the representative electrodes of the lobes. Then, we computed the distance of each electrode in the cap to the representative electrodes of each lobe and assigned it to one of the lobes if the distance between the given electrode and the closest lobe representatives was less than 2.3 cm. In this automatic algorithm, the electrodes located close enough to the representative electrodes of a lobe were assigned to that lobe. The topological plots in Fig. 1 show the distribution of the spatial locations of the electrodes assigned to each lobe according to the method based on the distance between electrodes and the representative electrodes of each lobe.

Following the method proposed by Liu and Falahpour, we used the band-specific EEG power measures to estimate a vigilance metric, as the ratio of the power in middle frequency bands (e.g., *α* and *β* bands) associated with increased wakefulness to the power in lower frequency bands (e.g., *δ* and *θ*) associated with decreased wakefulness (Klimesch, 1999; Liu and Falahpour, 2020). The higher this ratio, the higher the vigilance level. We subsequently used this vigilance measure to help explain inter-subject variations in the RVT-EEG associations.

### Cross Correlation Function Between EEG-bands Fluctuations and RVT

Since the correlation merely provides instantaneous zero-lag dependencies between two signals, we opted for the cross-correlation function (CCF) to characterize the dependency between EEG bands rhythms and RVT signals,

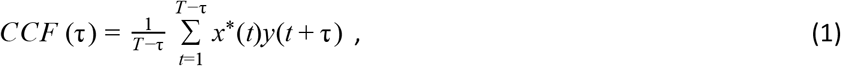

where *x(t)* and y(t) denote the band-specific EEG rhythm and RVT signal at time t, respectively. Since the hanning windows, applied for extracting PSD, implicitly result in smooth temporal fluctuations, we did not use further smoothing filters on each of them. Across subjects and lobes, these cross-correlation time series (CCF time series) exhibited temporal patterns in terms of peak time, strength, and more importantly the sign of correlations. In order to be flexible with respect to our hypotheses, we included a wide time window for the CCF computations, spanning a 100 s window (time lags of −50 s to 50 s).

### Statistical analysis

#### Surrogate data generation

Cross-correlation has been used in previous studies of a similar nature (Pfurtscheller et al., 2012; Yuan et al., 2013). In this work, the statistical significance of the peak of the cross correlation functions (CCFs) between the EEG-bands fluctuations and RVT were assessed via an iterative non-parametric approach, making 200 appropriate surrogates time series with the same probability distribution and the almost identical autocorrelations as the given time series (Schreiber and Schmitz, 1996). It is worth mentioning that the common alternative of using the amplitude adjusted Fourier transform (AAFT) algorithm proposed by Theiler et. al., in 1992 may lead to false positive statistical significance of the CCF peaks as the CCF analysis is highly sensitive to autocorrelations (Theiler et al., 1992).

#### Test for stationarity

To test for potential nonstationarities in the EEG and RVT data, we used the Kwaitkowski Phillips Schmidt Shin (KPSS) (Kwaitkowski et al., 1992) stationarity test in order to examine the RVT and EEG-bands fluctuation time series properties. The test of stationarity of the time series clearly shows that the use of methods for generating surrogate data that assume the stationarity of the EEG-bands rhythm and RVT may not lead to optimal results. Thus, we were prompted to additionally perform short-time Fourier transform and wavelet transform-based analyses.

#### Fourier & Wavelet Cross Spectrum Analysis (direction of correlation and non-stationarity)

Cross spectrum analyses are a class of powerful methods for investigating the synchronization of the cycles coming from two time series including period and phase. The simplest and most straightforward implementation of cross spectrum analysis is obtained by Fourier-transform the cross correlation of two signals. Fourier cross spectrum analysis can reveal rhythmic patterns that happen closely and strongly in two time series. However, the effectiveness of Fourier cross spectrum analysis is uncertain in the field of EEG and respiratory data due to the non-stationarity characteristics of them. Hence, we also provided wavelet cross spectrum analysis.

As has been shown in (Bigot et al., 2011), the wavelet cross spectrum analysis (WCS) provides a robust and consistent estimation of time-frequency relationship that may exist between two time data sets, especially in presence of non-stationary sources. Let consider X(*τ*,f) and Y(*τ*,f) as time-frequency representations of two given time series, the time-frequency based coherence between them at time τ and frequency f is defined as follows:

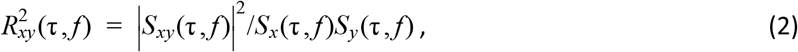

where *S_xy_*(*τ,f*) equals to *X*(*τ,f*)*Y*\*(*τ,f*), and *X*(*τ,f*) and *Y(τ,f)* are time-frequency representations of the signals (i.e., x(t) and y(t)), based either on the short-time Fourier transform of the wavelet transform. In the above definition, if *S_xy_(τ,f)* represents the wavelet cross spectrum (WCS) at frequency f and time τ between the time series x and y, the formula (2) will represent the definition of wavelet coherence between the given time series. In this study, we used the Morlet wavelet function, which has been widely used in the studies of EEG data to estimate the wavelet cross spectrum (Bigot et al., 2011). In the wavelet cross-coherence plot, the cone of influence represents the less accurate regions due to the edge effect at the beginning and end of the time series, and the phase represents the directionality of correlation between the two input signals.

**Figure 1.**
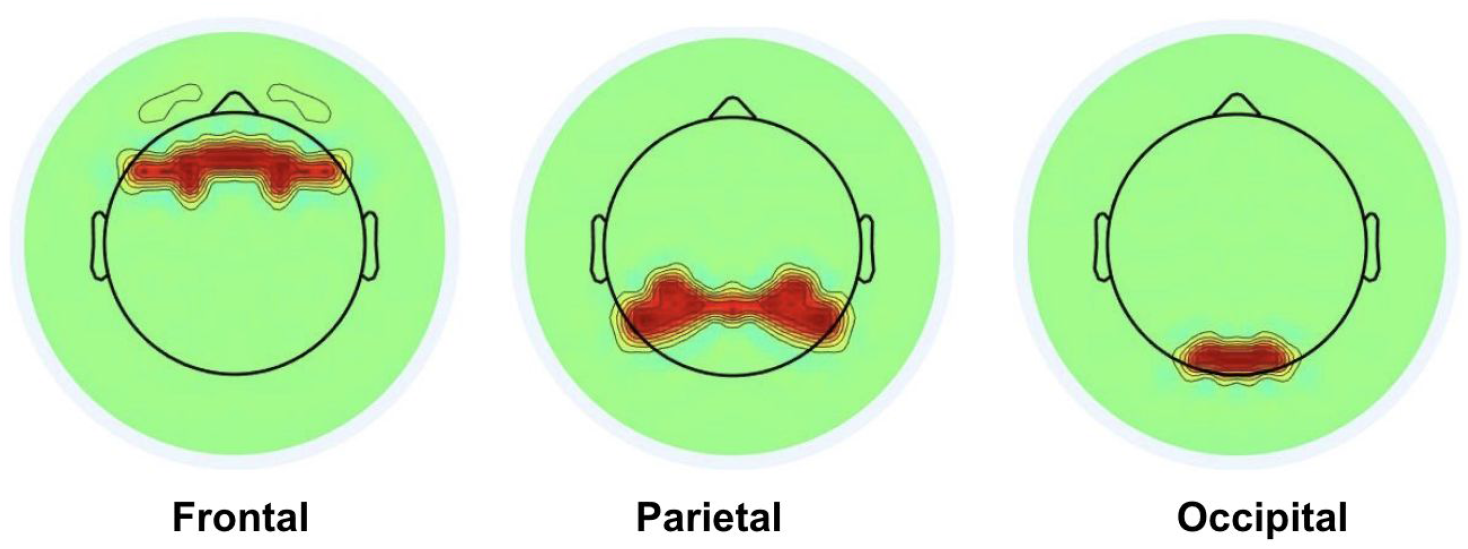
The topographic maps of the distribution of the spatial locations of the electrodes assigned to frontal, parietal, and occipital lobes resulted from the algorithm based on the distance of each electrode and the representative electrodes of the lobes. Since the topological mapping of the electrodes to the scalp view is a transition from low resolution to high resolution, a blurred coloured boundary is observed at the edges of the spatial distribution.

## Results

### Cross-correlation analysis

Shown in Fig. 2 are time series plots for a representative subject. Here we are demonstrating alpha, delta, theta and beta power spectrum fluctuations in a 150 s window. Also, the respiratory volume fluctuation (i.e., RVT) of the 150-second window for the representative subject is shown in Fig. 2.

**Figure 2.**
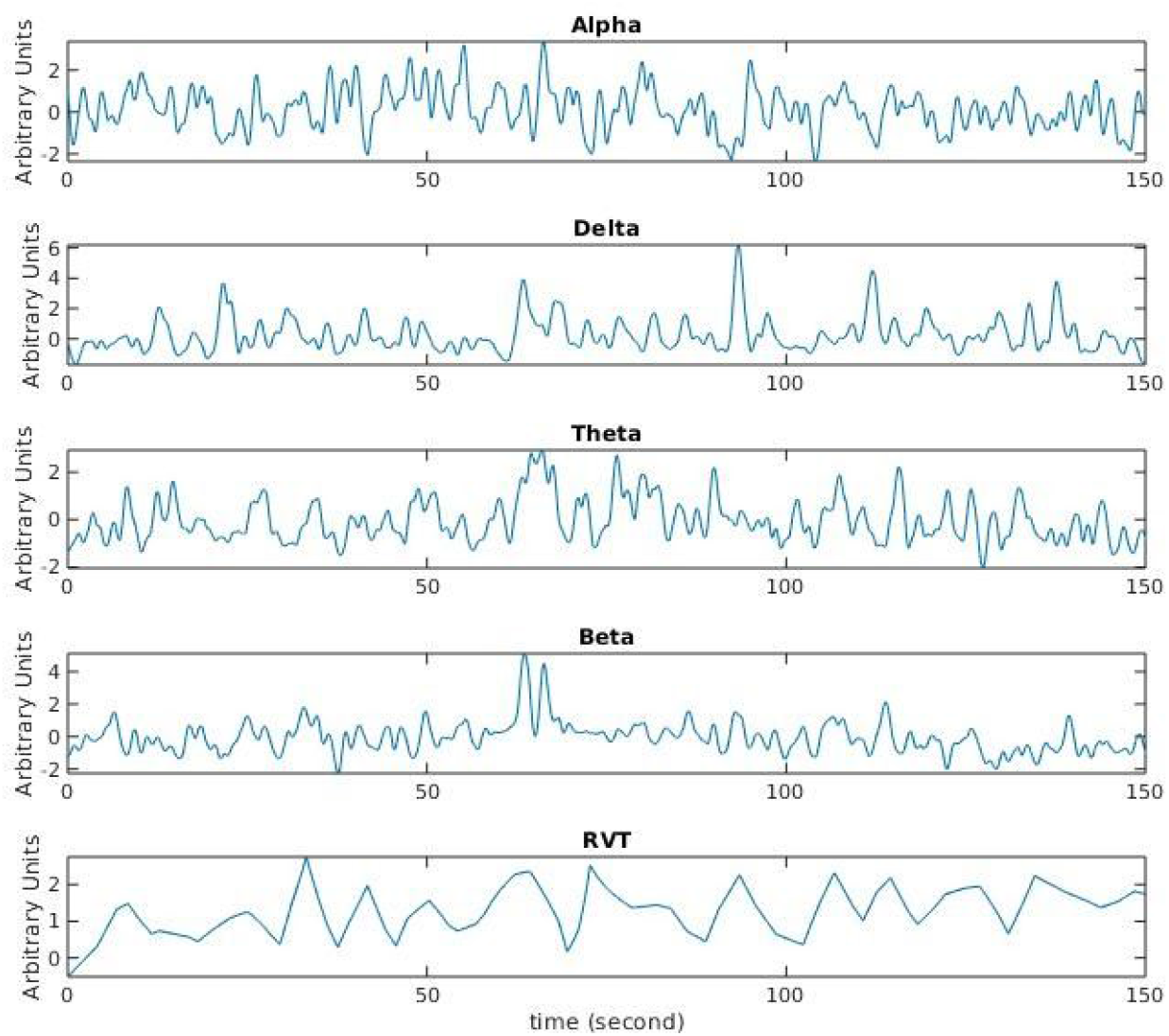
Sample EEG and RVT time series over a duration of 150 seconds, in the alpha, delta, theta and beta bands. Data are taken from a representative subject. The EEG time series are taken from the results of the PCA, and therefore have arbitrary units.

**Figure 3.**
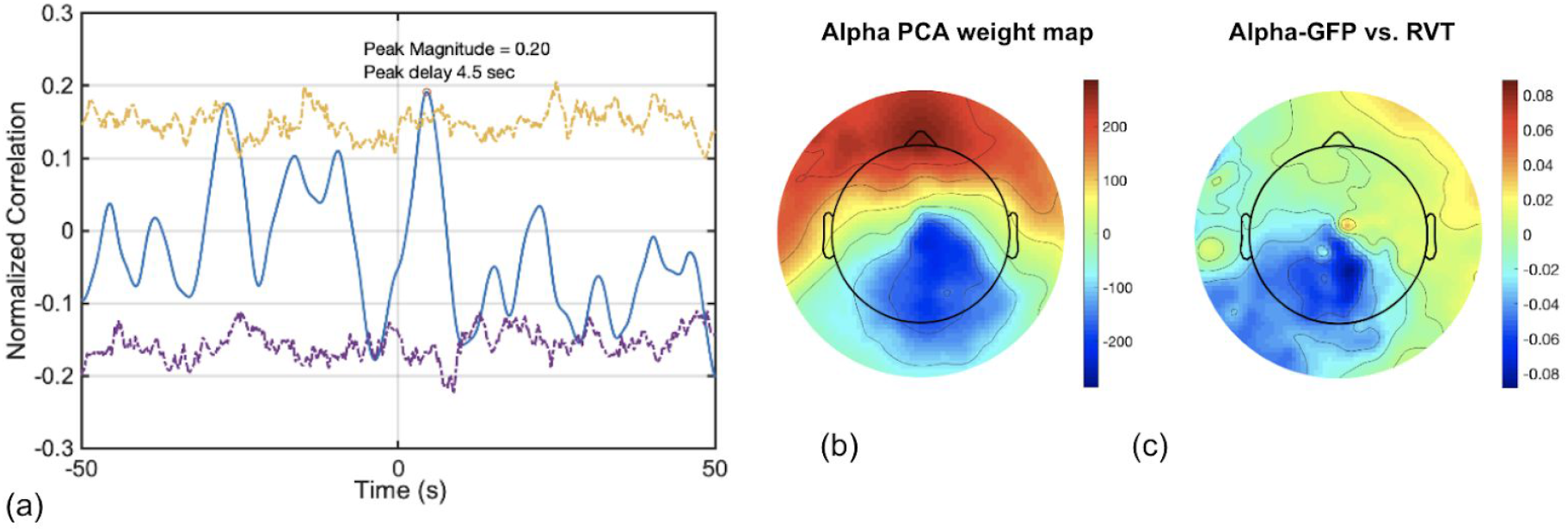
Sample cross-correlation plot for a representative subject. (a) Yellow and purple dashed lines respectively indicate the threshold of p =0.05 for highly positive and highly negative correlations at each lag, with the statistical significance determined using a set of surrogate time series. Statistically significant cross correlation peaks (blue) must surpass the boundaries marked by the dashed lines. (b) topological map of the PCA weights demonstrating the contribution of different EEG electrodes to the average alpha-GFP. (c) Topological map of the correlation between EEG time series and RVT at the RVT-GFP lag.

**Table 1.** Findings from cross-correlation analysis involving RVT and alpha power. For each subject, significant peaks within a 150 s window (at the p < 0.05 level, corrected for multiple comparisons) are shown in bold. In all cases, the associated time shifts (lags) are reported as well. Negative lags indicate that EEG leads RVT. Negative lags indicate that EEG leads RVT. Significant negative lags tend to be lower than significant positive lags, but the two are equally variable, as can be gleaned from the mean and STDEV for each. The mean and standard deviation of the peaks are lags are summarized for only the significant peaks.

| Subject | Frontal Lobe |  |  |  | Parietal Lobe |  |  |  | Occipital Lobe |  |  |  |
| --- | --- | --- | --- | --- | --- | --- | --- | --- | --- | --- | --- | --- |
|  | lag (s) | r | lag (s) | r | lag (s) | r | lag (s) | r | lag (s) | r | lag (s) | r |
| S1 | -1.6 | **0.21** | -7.2 | -0.21 | **0.8** | -0.1 | -2.9 | **0.2** | **2.2** | -0.17 | -2.6 | **0.22** |
| S2 | **7.7** | -0.19 | **12.7** | **0.14** | **8.00** | -0.22 | **12.9** | **0.12** | **8.6** | -0.13 | -2.10 | 0.11 |
| S3 | -2.00 | -0.07 | 5.20 | 0.07 | **12.0** | -0.2 | 2.50 | -0.14 | **2.3** | -0.22 | **7.7** | -0.26 |
| S4 | -0.5 | -0.11 | **15.9** | **0.14** | **15.9** | **0.2** | 16.00 | 0.17 | **16.1** | **0.12** | -10.60 | 0.12 |
| S5 | -9.70 | 0.05 | 14.60 | -0.08 | -2.40 | 0.07 | 9.50 | -0.08 | -2.3 | **0.11** | **10.6** | -0.11 |
| S6 | **0** | **0.23** | **6.7** | **0.19** | **0.0** | **0.3** | **5.2** | **0.3** | -9.70 | -0.16 | 5.10 | -0.08 |
| S7 | **4.2** | **0.12** | -4.6 | **0.14** | -4.40 | 0.12 | 8.90 | -0.11 | -4.6 | **0.12** | **8.8** | **0.10** |
| S8 | -2.8 | **0.22** | 7.40 | 0.25 | -3.4 | -0.3 | **6.8** | **0.2** | -3.4 | -0.27 | -8.40 | -0.23 |
| S9 | **15.2** | **0.16** | 0.20 | -0.10 | -1.8 | -0.1 | 15.10 | -0.09 | -1.00 | -0.13 | 1.80 | -0.11 |
| S10 | **0** | **0.42** | **6.9** | **0.17** | -10.5 | **0.1** | 7.20 | 0.10 | **3.7** | -0.11 | -0.30 | 0.10 |
| S11 | **1.3** | **0.38** | -4.6 | -0.16 | **1.6** | **0.4** | -4.8 | -0.2 | **1.5** | **0.33** | **12.7** | **0.30** |
| S12 | -1.0 | -0.18 | **7.3** | -0.28 | -3.90 | -0.08 | 3.00 | -0.09 | -2.00 | -0.09 | **13.90** | -0.12 |
| S13 | **3.9** | **0.22** | **9.6** | -0.14 | -0.9 | -0.1 | **5.1** | **0.2** | -6.10 | 0.13 | **11.00** | **0.19** |
| S14 | **1.5** | -0.14 | -5.8 | -0.23 | -8.5 | -0.2 | 10.70 | -0.14 | -8.7 | -0.21 | 5.00 | -0.14 |
| S15 | -12.90 | 0.12 | **2.70** | **0.23** | **9.0** | **0.2** | 11.90 | 0.16 | -2.6 | **0.17** | **10.4** | **0.19** |
| S16 | 7.70 | 0.11 | **10.50** | **0.11** | **14.7** | **0.2** | -4.20 | -0.10 | -10.00 | 0.17 | 10.40 | 0.19 |
| S17 | -10.20 | -0.22 | 4.60 | 0.09 | -3.1 | -0.2 | -13.90 | -0.15 | -13.00 | -0.21 | -7.60 | -0.21 |
|  | Negative lag |  | Positive lag |  | Negative lag |  | Positive lag |  | Negative lag |  | Positive lag |  |
| Mean | -4.3 | -0.060 | 7.4 | 0.077 | -4.5 | -0.10 | 7.9 | 0.12 | -4.1 | 0.021 | 8.1 | 0.0062 |
| STDEV | 3.2 | 0.19 | 4.9 | 0.20 | 3.3 | 0.16 | 5.1 | 0.21 | 2.3 | 0.19 | 4.5 | 0.20 |

**Table 2.** Findings from cross-correlation analysis involving RVT and delta power. For each subject, significant peaks within a 150 s window (at the p < 0.05 level, corrected for multiple comparisons) are shown in bold. In all cases, the associated time shifts (lags) are reported as well. Negative lags indicate that EEG leads RVT. As in the case of alpha, significant negative lags tend to be lower than significant positive lags, but the two are equally variable, as can be gleaned from the mean and STDEV for each. As well, there is a tendency for the negative r values to be associated with negative lags. The mean and standard deviation of the peaks are lags are summarized for only the significant peaks.

| Subject | Frontal Lobe |  |  |  | Parietal Lobe |  |  |  | Occipital Lobe |  |  |  |
| --- | --- | --- | --- | --- | --- | --- | --- | --- | --- | --- | --- | --- |
|  | lag (s) | r | lag (s) | r | lag (s) | r | lag (s) | r | lag (s) | r | lag (s) | r |
| S1 | **14.6** | **0.13** | -9.20 | -0.07 | **3.1** | **0.1** | **-3.5** | **0.15** | **-0.9** | **0.17** | **-13.7** | **-0.17** |
| S2 | 8.80 | -0.12 | -1.80 | 0.08 | **9.3** | **-0.11** | -5.80 | -0.13 | **10.2** | **-0.13** | -2.00 | 0.12 |
| S3 | **2.6** | **0.13** | 11.2 | 0.09 | -1.70 | 0.16 | 3.40 | 0.11 | 2.9 | 0.1105 | -6.40 | 0.11 |
| S4 | **3.8** | **-0.17** | 14.50 | 0.08 | **4.1** | **-0.15** | -13.60 | 0.08 | **4.20** | **-0.14** | -14.50 | 0.08 |
| S5 | **11.6** | **-0.19** | 2.10 | -0.16 | -9.90 | -0.05 | 8.00 | 0.10 | -3.70 | 0.06 | 8.60 | -0.11 |
| S6 | -1.90 | 0.15 | 12.80 | -0.14 | **-1.10** | **0.20** | 13.70 | -0.14 | -10.40 | -0.13 | 3.60 | -0.12 |
| S7 | **12.7** | **-0.15** | -4.00 | 0.13 | **12.4** | **-0.13** | -3.90 | 0.12 | 4.30 | 0.11 | 12.60 | -0.12 |
| S8 | **-2.8** | **-0.25** | **5.7** | **-0.3** | **1.9** | **0.28** | **-3.5** | **-0.26** | **2** | **-0.33** | **-4** | **-0.98** |
| S9 | **8.2** | **0.13** | -13.20 | -0.10 | **-0.4** | **-0.16** | -8.40 | -0.11 | **1.7** | **0.1** | **14.7** | **-0.14** |
| S10 | **0.3** | **0.39** | 9.30 | 0.08 | **-0.5** | **0.05** | - | - | **-0.9** | **0.56** | - | - |
| S11 | **1.6** | **0.19** | -3.20 | -0.14 | **0.3** | **0.27** | -13.20 | -0.16 | **1** | **0.18** | -13.40 | -0.14 |
| S12 | **0.3** | **0.14** | 9.30 | -0.08 | **5.5** | **-0.13** | - | - | -2.70 | 0.09 | **14.90** | **-0.14** |
| S13 | **10.5** | **-0.16** | -13.40 | 0.14 | **13.8** | **0.11** | -13.90 | -0.15 | **-4.7** | **-0.18** | **-8.5** | **0.16** |
| S14 | **-7.6** | **-0.18** | 6.80 | -0.13 | -7.50 | -0.10 | 6.10 | -0.09 | -7.10 | -0.13 | 6.00 | -0.12 |
| S15 | -2.30 | 0.14 | 13.70 | 0.13 | **2.5** | **0.22** | **8.6** | **0.14** | **2.7** | **0.17** | **14.9** | **-0.18** |
| S16 | **6.9** | **0.20** | 11.50 | 0.14 | **7.3** | **0.172** | **-3.5** | **-0.17** | **-3.00** | **-0.26** | 7.20 | 0.07 |
| S17 | **-12.1** | **-0.23** | -6.30 | -0.10 | **-12.2** | **-0.25** | 1.00 | -0.12 | **14.70** | **0.14** | **7.90** | **0.15** |
|  | Negative lag |  | Positive lag |  | Negative lag |  | Positive lag |  | Negative lag |  | Positive lag |  |
| Mean | -7.5 | -0.22 | 6.6 | 0.028 | -3.5 | -0.063 | 6.3 | 0.070 | -5.1 | -0.10 | 6.9 | -0.023 |
| STDEV | 4.7 | 0.04 | 5.0 | 0.21 | 4.1 | 0.19 | 4.4 | 0.17 | 4.6 | 0.48 | 5.9 | 0.20 |

**Table 3.**
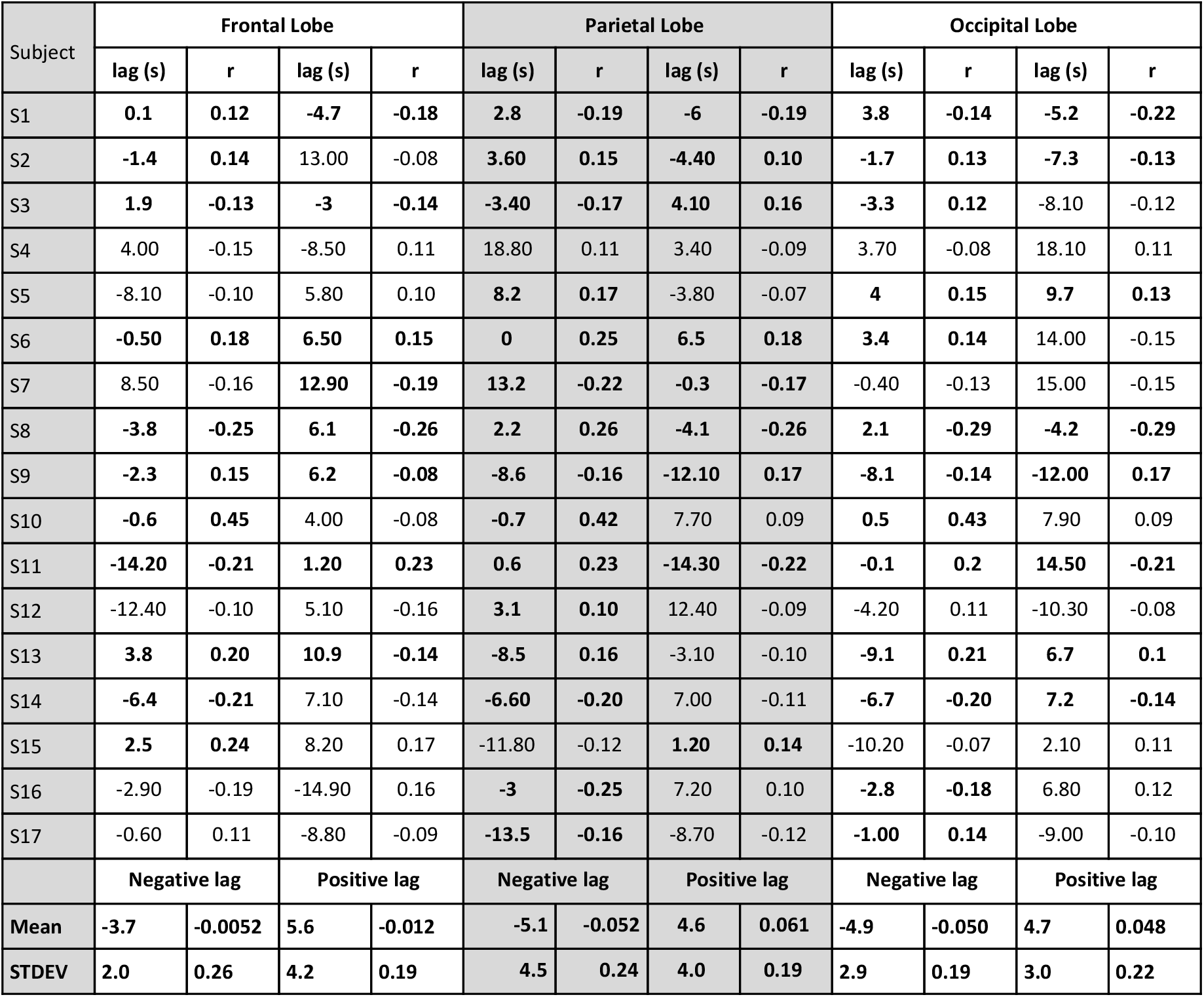
Findings from cross-correlation analysis involving RVT and theta power. For each subject, significant peaks within a 150 s window (at the p < 0.05 level, corrected for multiple comparisons) are shown in bold. In all cases, the associated time shifts (lags) are reported as well. Negative lags indicate that EEG leads RVT. Negative lags indicate that EEG leads RVT. In the theta case, significant negative lags are no longer lower than significant positive peaks, as was the case for both alpha and delta bands. As well, there is a tendency for the negative r values to be associated with negative lags. The mean and standard deviation of the peaks are lags are summarized for only the significant peaks.

| Subject | Frontal Lobe |  |  |  | Parietal Lobe |  |  |  | Occipital Lobe |  |  |  |
| --- | --- | --- | --- | --- | --- | --- | --- | --- | --- | --- | --- | --- |
|  | lag (s) | r | lag (s) | r | lag (s) | r | lag (s) | r | lag (s) | r | lag (s) | r |
| S1 | **0.1** | **0.12** | **-4.7** | **-0.18** | **2.8** | **-0.19** | **-6** | **-0.19** | **3.8** | **-0.14** | **-5.2** | **-0.22** |
| S2 | **-1.4** | **0.14** | 13.00 | -0.08 | **3.60** | **0.15** | **-4.40** | **0.10** | **-1.7** | **0.13** | **-7.3** | **-0.13** |
| S3 | **1.9** | **-0.13** | **-3** | **-0.14** | **-3.40** | **-0.17** | **4.10** | **0.16** | **-3.3** | **0.12** | -8.10 | -0.12 |
| S4 | 4.00 | -0.15 | -8.50 | 0.11 | 18.80 | 0.11 | 3.40 | -0.09 | 3.70 | -0.08 | 18.10 | 0.11 |
| S5 | -8.10 | -0.10 | 5.80 | 0.10 | **8.2** | **0.17** | -3.80 | -0.07 | **4** | **0.15** | **9.7** | **0.13** |
| S6 | **-0.50** | **0.18** | **6.50** | **0.15** | **0** | **0.25** | **6.5** | **0.18** | **3.4** | **0.14** | 14.00 | -0.15 |
| S7 | 8.50 | -0.16 | **12.90** | **-0.19** | **13.2** | **-0.22** | **-0.3** | **-0.17** | -0.40 | -0.13 | 15.00 | -0.15 |
| S8 | **-3.8** | **-0.25** | **6.1** | **-0.26** | **2.2** | **0.26** | **-4.1** | **-0.26** | **2.1** | **-0.29** | **-4.2** | **-0.29** |
| S9 | **-2.3** | **0.15** | **6.2** | **-0.08** | **-8.6** | **-0.16** | **-12.10** | **0.17** | **-8.1** | **-0.14** | **-12.00** | **0.17** |
| S10 | **-0.6** | **0.45** | 4.00 | -0.08 | **-0.7** | **0.42** | 7.70 | 0.09 | **0.5** | **0.43** | 7.90 | 0.09 |
| S11 | **-14.20** | **-0.21** | **1.20** | **0.23** | **0.6** | **0.23** | **-14.30** | **-0.22** | **-0.1** | **0.2** | **14.50** | **-0.21** |
| S12 | -12.40 | -0.10 | 5.10 | -0.16 | **3.1** | **0.10** | 12.40 | -0.09 | -4.20 | 0.11 | -10.30 | -0.08 |
| S13 | **3.8** | **0.20** | **10.9** | **-0.14** | **-8.5** | **0.16** | -3.10 | -0.10 | **-9.1** | **0.21** | **6.7** | **0.1** |
| S14 | **-6.4** | **-0.21** | 7.10 | -0.14 | **-6.60** | **-0.20** | 7.00 | -0.11 | **-6.7** | **-0.20** | **7.2** | **-0.14** |
| S15 | **2.5** | **0.24** | 8.20 | 0.17 | -11.80 | -0.12 | **1.20** | **0.14** | -10.20 | -0.07 | 2.10 | 0.11 |
| S16 | -2.90 | -0.19 | -14.90 | 0.16 | **-3** | **-0.25** | 7.20 | 0.10 | **-2.8** | **-0.18** | 6.80 | 0.12 |
| S17 | -0.60 | 0.11 | -8.80 | -0.09 | **-13.5** | **-0.16** | -8.70 | -0.12 | **-1.00** | **0.14** | -9.00 | -0.10 |
|  | Negative lag |  | Positive lag |  | Negative lag |  | Positive lag |  | Negative lag |  | Positive lag |  |
| Mean | -3.7 | -0.0052 | 5.6 | -0.012 | -5.1 | -0.052 | 4.6 | 0.061 | -4.9 | -0.050 | 4.7 | 0.048 |
| STDEV | 2.0 | 0.26 | 4.2 | 0.19 | 4.5 | 0.24 | 4.0 | 0.19 | 2.9 | 0.19 | 3.0 | 0.22 |

**Table 4.** Findings from cross-correlation analysis involving RVT and beta power. For each subject, significant peaks within a 150 s window (at the p < 0.05 level, corrected for multiple comparisons) are shown in bold. In all cases, the associated time shifts (lags) are reported as well. Negative lags indicate that EEG leads RVT. Negative lags indicate that EEG leads RVT. In the beta case, the occipital lobe is associated with fewer significant r values, significant negative lags are no longer lower than significant positive peaks, as was the case for both alpha and delta bands. As well, there is no sign that the negative r values are more often associated with negative lags. The mean and standard deviation of the peaks are lags are summarized for only the significant peaks.

| Subject | Frontal Lobe |  |  |  | Parietal Lobe |  |  |  | Occipital Lobe |  |  |  |
| --- | --- | --- | --- | --- | --- | --- | --- | --- | --- | --- | --- | --- |
|  | lag (s) | r | lag (s) | r | lag (s) | r | lag (s) | r | lag (s) | r | lag (s) | r |
| S1 | **-13.0** | **0.1** | 4.8 | 0.07 | **-1.4** | **0.22** | **-5.7** | **-0.16** | **-1.2** | **0.25** | **2.8** | **0.2** |
| S2 | **-2.80** | **0.27** | **5.6** | **0.16** | **2.2** | **-0.32** | **-2.2** | **0.3** | **2.2** | **-0.3** | **-2.6** | **0.24** |
| S3 | **1.30** | **-0.14** | -3.5 | -0.09 | **1.3** | **-0.13** | **-3.8** | **-0.15** | **-3.4** | **0.13** | 6.2 | 0.1 |
| S4 | **3.20** | **0.15** | -7.0 | 0.1 | **4.0** | **-0.17** | -6.5 | 0.1 | **4.0** | **-0.17** | -6.5 | 0.1 |
| S5 | -4.2 | -0.1 | 1.7 | 0.09 | -5.1 | 0.0 | 1.1 | 0.07 | **-0.8** | **0.1** | 6.6 | -0.1 |
| S6 | **0.70** | **0.23** | **8.6** | **0.22** | **0.10** | **0.28** | **7.3** | **0.19** | **-10.2** | **0.15** | 11.8 | 0.1 |
| S7 | **-5.7** | **0.14** | 4.7 | 0.12 | **-4.5** | **0.15** | 14.9 | -0.10 | **-5.7** | **0.1** | 8.8 | -0.1 |
| S8 | **-2.6** | **-0.32** | **4.6** | **-0.3** | **-7.2** | **-0.3** | 4.6 | 0.21 | **3.2** | **-0.34** | - | - |
| S9 | **-7.1** | **-0.18** | 13.2 | 0.12 | **-7.3** | **-0.2** | -12.2 | 0.13 | -0.9 | -0.1 | -7.6 | -0.1 |
| S10 | **-0.4** | **0.52** | 6.9 | 0.12 | **-0.40** | **0.36** | 5.6 | -0.09 | **5.9** | **-0.13** | -12.5 | 0.1 |
| S11 | **0.10** | **0.42** | 14.8 | 0.1 | **0.40** | **0.43** | -6.0 | -0.1 | **0.60** | **0.40** | **-12.4** | **-0.2** |
| S12 | **5.9** | **-0.18** | -11.9 | -0.1 | **5.1** | **0.16** | -11.8 | -0.1 | -10.7 | -0.1 | 7.1 | -0.1 |
| S13 | **-5.1** | **-0.16** | -0.8 | 0.11 | **-8.9** | **0.15** | -4.6 | -0.13 | **-8.3** | **0.19** | -4.3 | -0.1 |
| S14 | **7.7** | **-0.2** | **-7.3** | **-0.23** | **-6.7** | **-0.24** | 6.6 | -0.17 | -6.4 | -0.2 | 3.2 | -0.1 |
| S15 | **3.7** | **-0.16** | **8.5** | **0.19** | -13.4 | -0.1 | 5.0 | 0.13 | 13.9 | 0.1 | 3.5 | 0.1 |
| S16 | **11.5** | **0.24** | -5.6 | -0.1 | 2.3 | 0.1 | 8.1 | 0.09 | -5.0 | -0.1 | 12.1 | 0.1 |
| S17 | -8.8 | -0.2 | 15.4 | -0.11 | **1.7** | **-0.15** | **-7** | **-0.14** | **1.8** | **-0.13** | **-5.4** | **-0.15** |
|  | Negative lag |  | Positive lag |  | Negative lag |  | Positive lag |  | Negative lag |  | Positive lag |  |
| Mean | -5.5 | 0.02 | 5.1 | 0.04 | -4.8 | 0.03 | 2.7 | 0.07 | -5.2 | 0.14 | 2.9 | -0.07 |
| STDEV | 3.9 | 0.29 | 3.7 | 0.24 | 2.8 | 0.23 | 2.4 | 0.27 | 3.5 | 0.15 | 1.7 | 0.27 |

As can be seen in Tables 1–4, there are statistically significant cross-correlations identified between RVT and EEG in all lobes and all frequency bands investigated, and these were observed for nearly all subjects. Given the complexity of respiration-mediated autonomic function, we did not constrain the direction of the cross correlation, thus allowing the investigation of either EEG leading RVT or vice versa. In this case, negative time lags indicate that EEG leads RVT.

For alpha LSFP (Table 1), significant negative lags tend to be lower than significant positive lags, but the two are equally variable, as can be gleaned from the mean and STDEV for each. As well, there is a tendency for the negative r values to be associated with negative lags. Moreover, most of the subjects exhibited > 1 significant correlation peak within the 100-sec window. The histogram of lags and r values associated with significant alpha-RVT CCF peaks are shown in Fig. 4a. The peak r values are distributed around ± 0.2, with the corresponding time lags distributed between 0 and ~15 s. Alpha LSFP leads RVT by 4.3 s, 4.5 s and 4.1 s in the frontal, parietal and occipital lobes, respectively. In cases whereby RVT leads alpha LSFP, the lags are 7.4 s, 7.9 s and 8.1 s in the three lobes, respectively. There is a preponderance of peaks associated with negative lags (EEG leading RVT) in the frontal lobe, but this is not observed in the parietal and occipital lobes.

**Figure 4.**
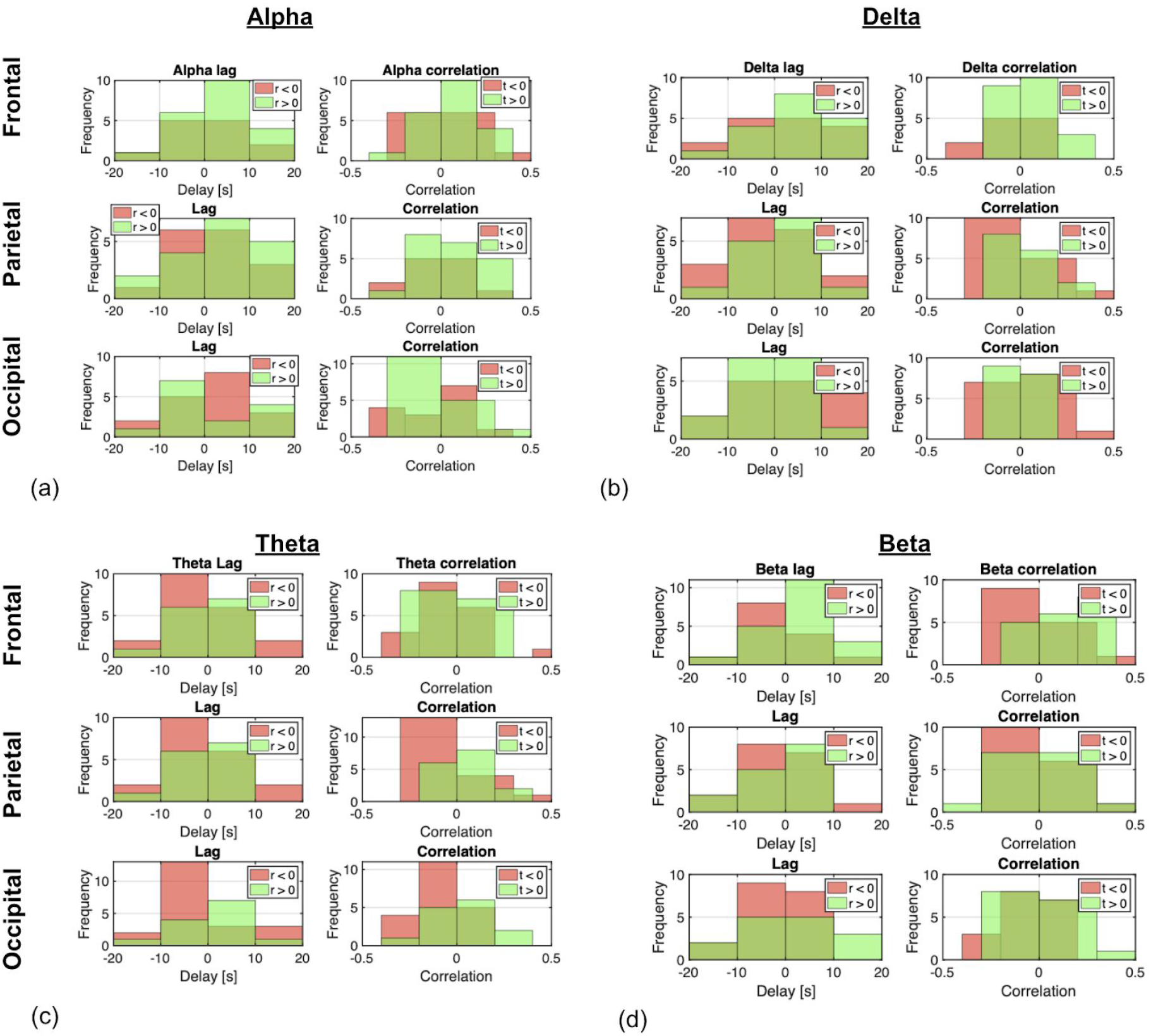
Distribution of the peak cross-correlation coefficients and their corresponding time lags for each frequency band, shown for the frontal, parietal and occipital lobes. These plots summarize the results in Tables 1–4, and the data are divided into categories that depend on the polarity of the lags (t) and associated correlations (r). The peak r values are distributed around ± 0.2, with the corresponding time lags distributed between 0 and ~15 s. The lag and r value histograms are colour-coded with the signs of the respective r values and lags, as indicated in the legend. The darker green indicates overlaps between the two categories. All CCF peaks from Tables 1–4 are included in this plot. Visually, negative peak correlations tend to be more often associated with negative lags in all four bands and all 3 lobes. More specifically, this effect is the most dominant in the theta band, followed by the remaining bands. This effect is also less observed in the frontal lobe compared to the parietal and occipital lobes. In the occipital lobe, compared to other lobes, the alpha wave is associated with a higher number of negative correlations at positive lags (EEG lagging RVT). Note: Negative lags indicate that EEG leads RVT.

In the delta band (Table 2), as in the case of alpha, significant negative lags tend to be lower than significant positive lags, but the two are equally variable, as can be gleaned from the mean and STDEV for each. As well, there is a tendency for the negative r values to be associated with negative lags. However, unlike in the alpha band, fewer subjects exhibit multiple significant CCF peaks. In cases of negative CCF peaks (delta LSFP leads RVT), the lags are 7.5 s, 3.5 s and 5.1 s in the frontal, parietal and occipital lobes, respectively. In cases whereby RVT leads delta LSFP, the lags are 6.6 s, 6.3 s and 6.9 s in the three lobes, respectively.

On the other hand, in the theta band (Table 3), significant negative lags are no longer lower than significant positive peaks, as was the case for both alpha and delta bands. As well, there is a tendency for the negative r values to be associated with negative lags. Theta LSFP leads RVT by 3.7 s, 5.1 s and 4.9 s in the frontal, parietal and occipital lobes, respectively. In cases whereby RVT leads theta LSFP, the lags are 5.6 s, 4.6 s and 4.7 s in the three lobes, respectively.

Lastly, in the beta band (Table 4), the occipital lobe is associated with fewer significant r values, significant negative lags are no longer lower than significant positive peaks, as was the case for both alpha and delta bands. Beta LSFP leads RVT by 5.5 s, 4.8 s and 5.2 s in the frontal, parietal and occipital lobes, respectively. In cases whereby RVT leads alpha LSFP, the lags are 5.1 s, 2.7 s and 2.9 s in the three lobes, respectively. There is no sign that the negative r values are more often associated with negative lags.

As illustrated in Fig. 4, there is no preponderance of negative lags (EEG leading RVT) over positive lags. The peak r values are distributed around ± 0.2, associated with lags that are mainly distributed between 0 and ± 10 s. One generalizable observation, however, is that negative peak correlations tend to be more often associated with negative lags in all four bands and all 3 lobes.

**Table 5.** Summary of statistically significant peak-lag times for different EEG bands and lobes.

| Lobe | EEG leading RVT |  |  |  | Mean |
| --- | --- | --- | --- | --- | --- |
|  | Delta | Theta | Alpha | Beta |  |
| Frontal | 7.5 | 3.7 | 4.3 | 5.5 | 5.3 ±1.5 |
| Parietal | 3.5 | 5.1 | 4.5 | 4.8 | 4.5 ±0.6 |
| Occipital | 5.1 | 4.9 | 4.1 | 5.2 | 4.8 ±0.4 |
| Mean | 5.2±1.6 | 4.6±0.6 | 4.3±0.2 | 5.2±0.3 | |

| Lobe | RVT leading EEG |  |  |  | Mean |
| --- | --- | --- | --- | --- | --- |
|  | Delta | Theta | Alpha | Beta |  |
| Frontal | 6.6 | 5.6 | 7.4 | 5.1 | 6.6 ±0.9 |
| Parietal | 6.3 | 4.6 | 7.9 | 2.7 | 5.4±1.9 |
| Occipital | 6.9 | 4.7 | 8.1 | 2.9 | 5.7±2.0 |
| Mean | 6.6±0.2 | 5.0±0.4 | 7.8±0.3 | 3.6±1.1 |  |

**Figure 5.**
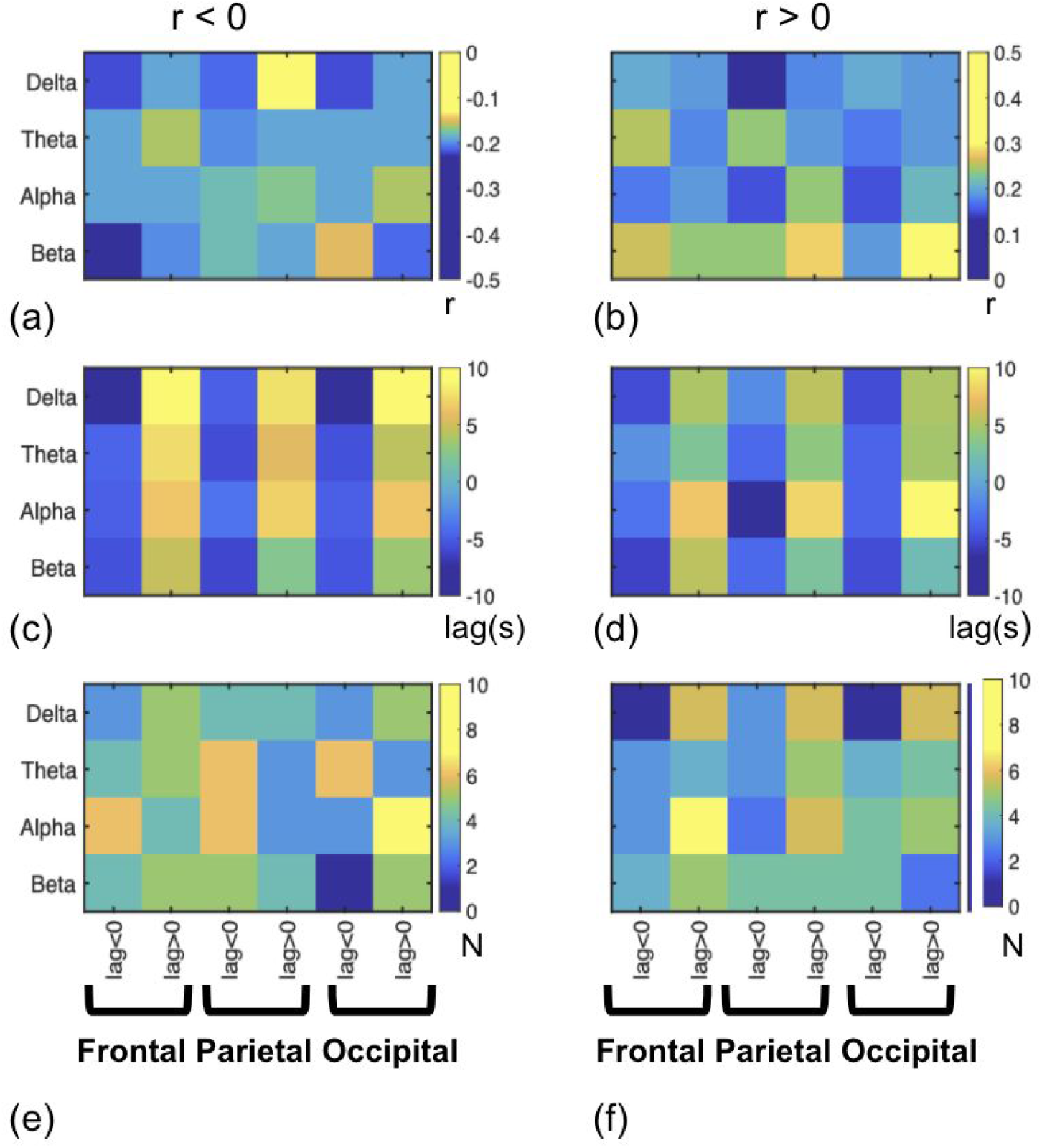
Summary of statistically significant peak correlation (r), the associated lags and number of subjects displaying each scenario (N). (a), (c), (e) correspond to cases in which peak r > 0, and (b), (d) and (f) correspond to the opposite. In each matrix, alternating columns reflect positive and negative lags. Negative lags indicate that EEG leads RVT.

In Fig. 5 it can be observed that the delta band is typically associated with larger lag values for r < 0 (Fig. 5c), and whereas alpha band is associated with larger lags for r > 0 (Fig. 5d). Also, more subjects appear to be associated with r < 0 than with r > 0 in the theta and alpha bands (Fig. 5e & f). Nonetheless, there is no visible difference across lobes.

**Figure 6.**
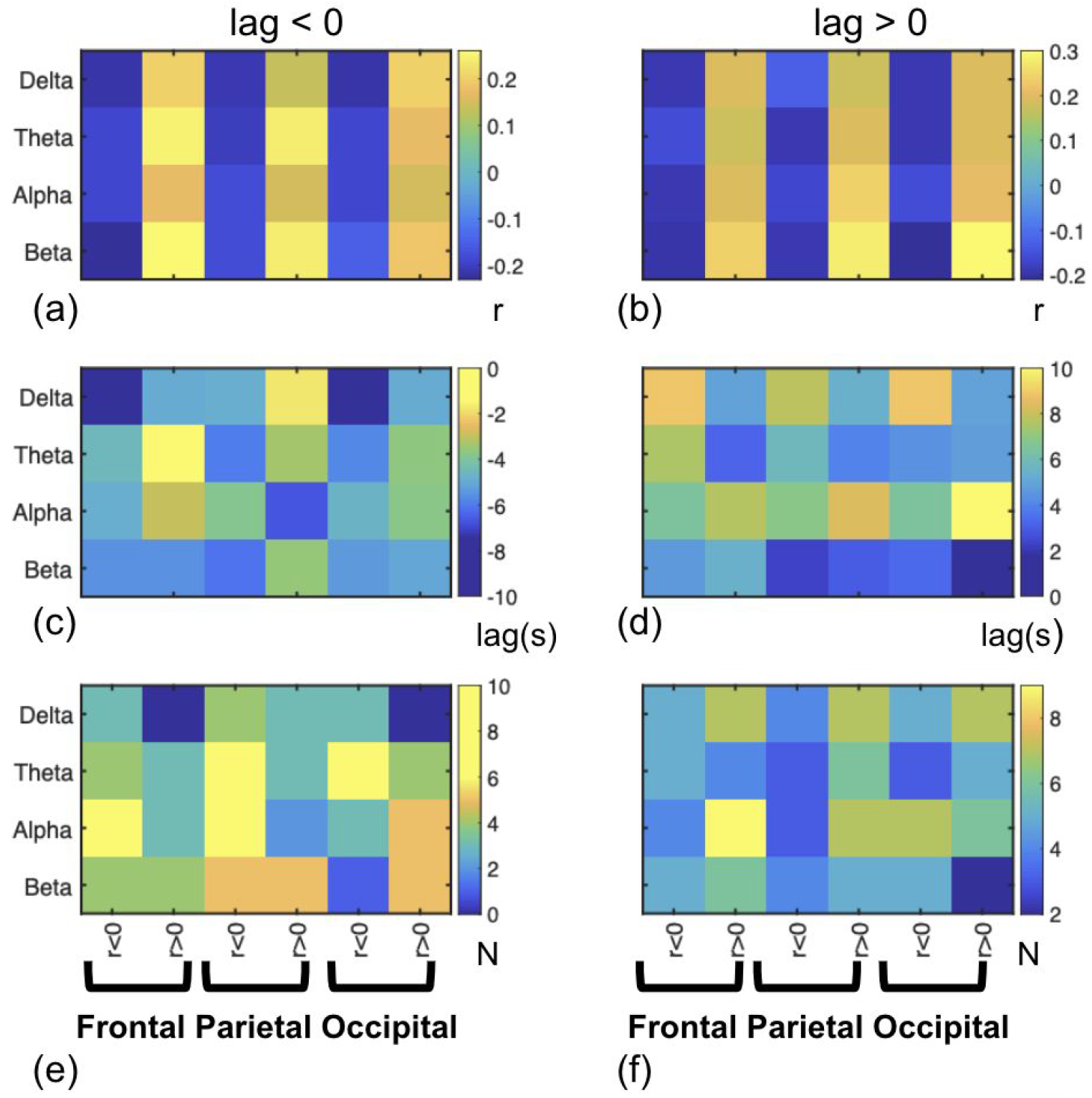
Summary of statistically significant peak correlation (r), the associated lags and number of subjects displaying each scenario (N). (a), (c), (e) correspond to cases in which lag > 0, and (b), (d) and (f) correspond to the opposite. Negative lags indicate that EEG leads RVT. In each matrix, alternating columns reflect positive and negative r values.

In Fig. 6 it can be seen that the delta band is associated with the largest negative lags (Fig. 6c), whereas the alpha band is associated with the largest positive lags (Fig. 6d). Moreover, more individuals exhibit negative than positive lags (e). As well, the theta and alpha band are associated with higher positive r values when lag < 0 (Fig. 6a). Also, the parietal lobe is associated with the highest number of subjects exhibiting a negative lag (EEG leads RVT) (Fig. 6e), as well as a larger number of subjects exhibiting positive lag (EEG lags RVT) in the frontal lobe (Fig. 6f).

**Figure 7.**
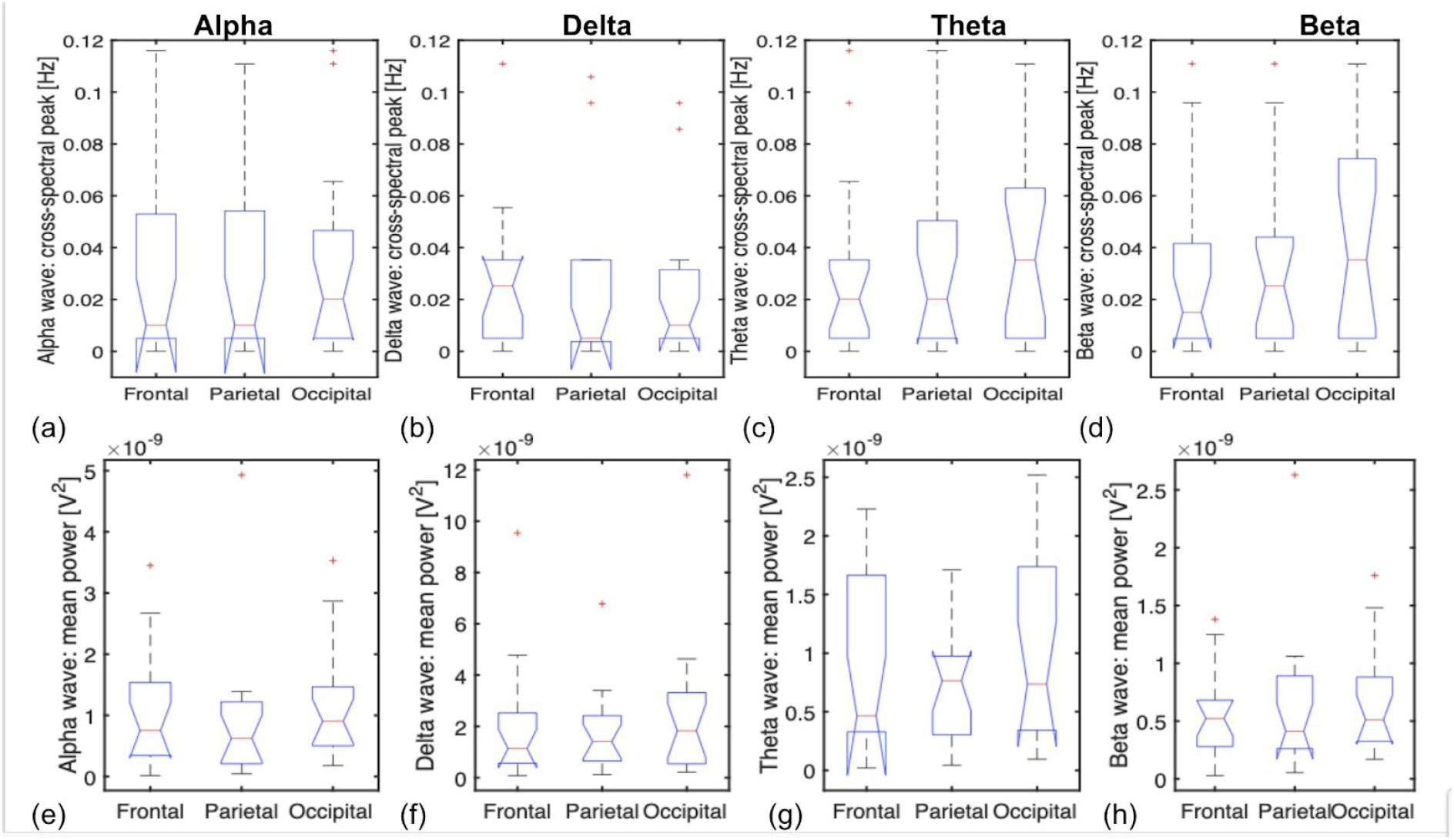
Cross-spectral peak frequencies and EEG LSFP amplitudes (different lobes and bands). (a-d) It can be observed across multiple brain regions, that the alpha-RVT and delta-RVT associations on average occur at ≤0.02 Hz (indicated by the red lines), whereas the theta-RVT and beta-RVT associations appear to peak at ≥0.02 Hz. (e-h) Moreover, the inter-lobe variations in band-specific EEG power are unrelated to the inter-lobe variations in the location of the peak coherence.

Given the large inter-subject variability in the EEG-RVT relationships in all lobes and frequency bands, we further probed whether this variability is associated with variability in vigilance across subjects. We failed to observe such an association except in the case of occipital beta LFSP, whereby the peak cross-correlation r value is significantly and positively correlated with vigilance (Fig. 8).

**Figure 8.**
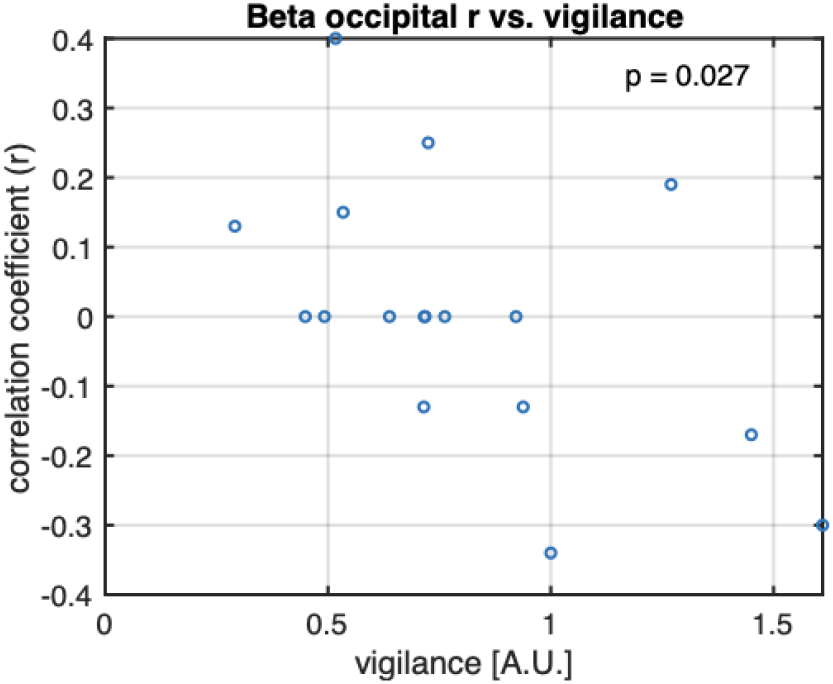
The peak cross-correlation coefficient (r) as a function of vigilance. The only significant association was found in occipital beta power.

### Wavelet cross-coherence

The use of wavelet cross-coherence (WCC) was motivated by the results of the stationarity tests, which demonstrated that both EEG (alpha GFP) and RVT signals are nonstationary in the majority of our subjects (Table A1 in the Supplementary materials). The WCC analyses revealed that the relationship between EEG LPSF and RVT is only intermittently in-phase. This in-phase relationship is mainly found in the range of 0.03 - 0.1 Hz. This finding is representative of the rest of our subjects, as can be corroborated by the data in Fig. 7(a-d). These figures demonstrate that the relationship between EEG and RVT is non-stationary. The colour bar represents the coherence values, with yellow indicating high coherence.

**Figure 9.**
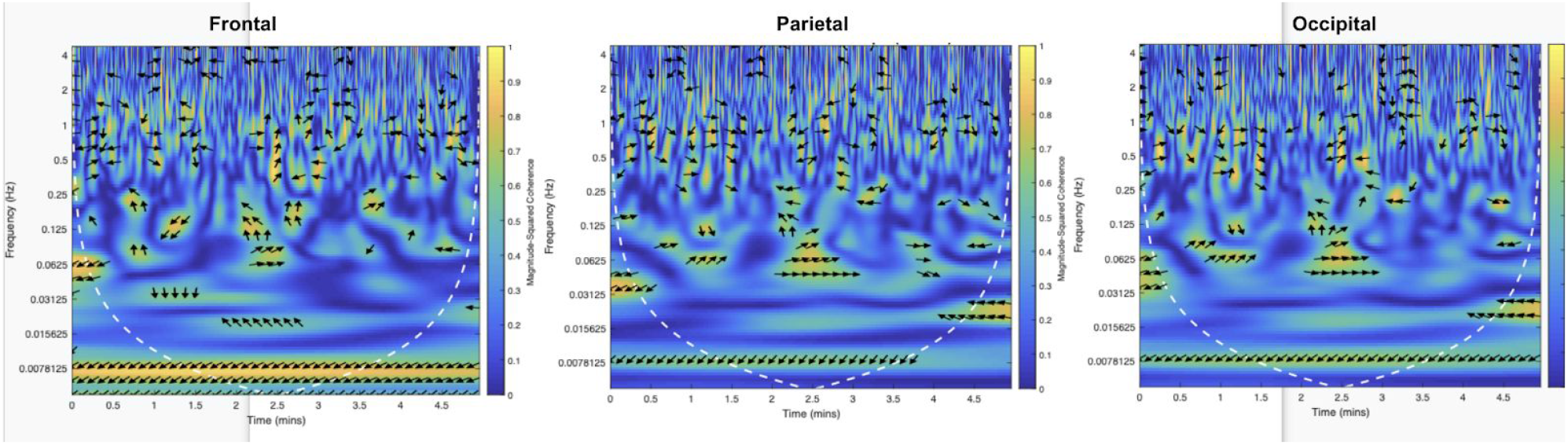
Wavelet coherence for alpha LSFP and RVT, for one representative subject. The y axis represents the frequency, and the x axis time. The dashed white line indicates the cone of influence, where edge effects occur in the coherence data. The arrows represent the phase relationship between EEG and RVT, with rightward arrows indicating an intermittent in-phase relationship between alpha LSFP and RVT. The in-phase relationship is mainly found in the range of 0.03 - 0.1 Hz. This finding is representative of the rest of our subjects. These figures demonstrate that the relationship between EEG and RVT is non-stationary. The colour bar represents the coherence values, with yellow indicating high coherence.

## Discussion

In the resting-state fMRI (rs-fMRI) literature, RVT is generally considered to be a low-frequency physiological nuisance. However, the lack of neuronal association with RVT has yet to be proven. Due to the close association between respiration, heart rate and autonomic nervous regulation (Aguirre et al., 1990; Iacovella and Hasson, 2011), and in turn, the association between the latter and cognition, it is reasonable to expect RVT to have neuronal associations. In fact, a recent study found that the alpha-EEG-RVT association is arousal-dependent (Yuan et al., 2013), leading us to question the participation of other EEG bands (hence domains of brain function) in the EEG-RVT connection. Moreover, while the association between RVT and the rs-fMRI signal has been reproduced in several studies (Birn et al., 2006; Chang and Glover, 2009; Falahpour et al., 2013; Golestani et al., 2015), we seek the neural component of fMRI alone. Thus, we bypass rs-fMRI in this study and focus on investigating the relationship RVT and EEG in detail, in the hope of establishing whether there is a generalizable association between EEG and RVT, across multiple frequency bands and brain regions.

### Basis of association between respiratory and neural activity

The neuronal involvement in breathing tasks has been well studied over the years, since it was first demonstrated in hedgehogs (Adrian, 1942). It is well known that respiratory control resides in the brainstem (Pattinson et al., 2009). However, respiratory changes have been associated with functional involvement in cortical and subcortical regions (Evans, 2010). McKay et al. reported using fMRI that changes in respiration pattern were associated with significant alterations in neuronal activity in the motor-sensory cortices, supplementary motor area, thalamus, caudate, globus pallidum and the cerebellum (McKay et al., 2003). Moreover, breathholding was found to be associated with fMRI signal changes in the the insula, basal ganglia, frontal cortex, parietal cortex and thalamus, which are in common with response inhibition tasks, and in addition, activity within the pons (McKay et al., 2008).

In terms of EEG-based measurements, Busek and Kemlink reported an increase in delta power in the anterior temporal cortex during slow breathing (breathing frequency = 0.1 - 0.25 Hz) compared to regular-paced breathing, as well as a reduction in both delta and theta power when breath-rate reached 0.5 Hz (Busek and Kemlink, 2005). Furthermore, meditation, often involving slowing of breathing, is shown to increase EEG power in the alpha, beta and theta bands, being more pronounced in the temporal and occipital regions in the case of alpha (Ahani et al., 2014). Hudson et al. observed increase in EEG power in the 0-5 Hz range due to breathing tasks, but no significant EEG associations in resting-state spontaneous breathing (Hudson et al., 2016). Heck et al. further proposed that respiration connects with high cognitive functions to form a part of ongoing cortical rhythmic activity (Heck et al., 2016).

As RVT is naturally associated with fluctuations in end-tidal carbon dioxide (PETCO2), albeit nonlinearly (Morelli et al., 2018), arterial CO2 is a common target in biofeedback training. Biofeedback involves the autonomic nervous system (ANS), in which elevations in arterial CO2 are associated with increased drowsiness (Naifeh et al., 1982). As part of ANS regulation, the prefrontal and cingulate cortex exert tonic inhibitory control over the brainstem (Lane et al., 2009). Furthermore, sensory integration of respiration has also involved an interplay of volitional respiratory control (premotor and supplementary motor area (SMA)) and interoceptive integration areas (anterior cingulate cortex, insula, and amygdala) (Evans, 2010). Furthermore, it has been demonstrated that consciously controlled change in respiratory behavior will cause a change in cognitive and emotional states (Heck et al., 2016). Slowed breathing leads to a shift of ANS regulation from sympathetic (SNS) to parasympathetic (PNS) (Jerath et al., 2006), which is in turn related to a decrease in heart rate and increased high-frequency heart-rate variability. The increase in PNS regulation has been associated with better cognitive performance (Matthews, 2004) and great emotional response (O’Connor et al., 2007). However, it is unclear whether these cognitive connections of respiration extend to the resting state, in which respiratory variations are likely unconsciously driven.

### Resting-state respiration and neural activity: Cross correlations

As mentioned previously, despite the consensus that RVT constitutes a noise source in rs-fMRI (Birn et al., 2006; Chang and Glover, 2009; Falahpour et al., 2013), the possible neural associations of RVT have been under-investigated. Past investigations into the associations of EEG with spontaneous breathing have mostly focused on the alpha band. Fumoto et al. (Fumoto et al., 2004) observed an increase in high-frequency alpha power (10-13 Hz) during abdominal breathing, one that is distinct in frequency to the alpha increase in the eyes-closed state, and one which was attributed to an increase in serotonin and the action of serotonergic neurons. Most recently, in the work of Keller et al., respiratory (and heart-rate) fluctuations in the intermediate-frequency range (centred at 0.15 Hz) correlated with the fMRI signal in the mid and posterior insula and the secondary somatosensory area, which are regions related to interoceptive perception (Keller et al., 2020); it was proposed that respiratory variability shares a common central-nervous pathway with heart-rate variability, and that low-frequency respiratory rhythms can in fact modulate higher-frequency EEG oscillations through a harmonics relationship (Mather and Thayer, 2018). Specifically in the context of rs-fMRI, there has only been a single study to date; Yuan et al. demonstrated that alpha global-field power (GFP) is significantly associated with RVT in the eyes-closed state (Yuan et al., 2013), with alpha-GFP leading RVT by 5.20 s, at a r value of 0.4.

In this work, we began with the approach of Yuan et al. (Yuan et al., 2013). That is, we time shifted RVT and EEG power with respect to each other to examine their correlations. We found that RVT is significantly associated with EEG lobe-specific field power (LSFP) in all bands investigated, but that the EEG-RVT time lag varies with EEG band and brain region (Table 5). We included both positive and negative CCF peaks, with both positive and negative lags. Specifically, we found that in the eyes closed condition, alpha LSFP leads RVT by 4.3 s, 4.5 s and 4.1 s in the frontal, parietal and occipital lobes, respectively. These values are in line with those reported by Yuan et al, and can be interpreted as a sign of ANS regulation leading to downstream breathing changes. The lag between brainstem signalling and detectable changes in respiration is well within the range of normal lags in respiratory regulation (Ben-Tal and Smith, 2010). Conversely, there are also a sizable number of cases in which RVT leads alpha LSFP, with lags of 7.4 s, 7.9 s and 8.1 s in the three lobes, respectively. On average, the distance between the positive and negative CCF peaks are approximately 12 s (similar across all lobes), which, if representative of a pseudo-periodic wavelength, translates to 0.08 Hz.

For the delta band, in cases of negative CCF peaks (LSFP leads RVT), the lags are 7.5 s, 3.5 s and 5.1 s in the frontal, parietal and occipital lobes, respectively. These lags demonstrate a higher inter-regional variability than in the case of alpha LSFP. As discussed earlier, delta power is expected to exhibit a frontal prominence. In cases whereby RVT leads delta LSFP, the lags are 6.6 s, 6.3 s and 6.9 s in the three lobes, respectively. These values are aligned with (though slightly lower than) those from the alpha LSFP. On average, the distance between the positive and negative CCF peaks range between 9 s and 14 s, depending on the lobe. Nevertheless, in anesthetized rats, the feedback mechanism appears to be absent (Musizza et al., 2007), suggesting that the delta-RVT relationship may be arousal-dependent.

For the theta band, when LSFP leads RVT, the lags are 3.7 s, 5.1 s and 4.9 s in the frontal, parietal and occipital lobes, respectively. These are shorter than the lags associated with the delta band, particularly in the frontal lobe. In cases whereby RVT leads theta LSFP, the lags are 5.6 s, 4.6 s and 4.7 s in the three lobes, respectively. These lags are in the same range as those found for alpha and delta LSFPs. The average interpeak distance ranges between 9 s and 10 s, rather shorter than for the other bands. The theta band is known to be more prominent in the awake state. Our results are supported by evidence from the mouse brain supporting widespread coupling between theta activity and respiration (Tort et al., 2018), with neural field potentials entraining respiration.

Lastly, the beta band, which is of higher frequency than the three other bands, exhibited less robust LSFP-RVT associations than the other 3 bands (particularly in the occipital lobe). This is to be expected, given the higher noise content in higher-frequency EEG bands. The weaker beta associations are expected, as beta is lower in the eyes-closed state, which our subjects were in. Nonetheless, over half of all subjects exhibited significant associations. Beta LSFP leads RVT by 5.5 s, 4.8 s and 5.2 s in the frontal, parietal and occipital lobes, respectively. In cases whereby RVT leads alpha LSFP, the lags are 5.1 s, 2.7 s and 2.9 s in the three lobes, respectively. Beta power increase has been associated with meditation in which alternate nostril breathing was performed (Stancák and Kuna, 1994), but this phenomenon implicates a respiration-locked EEG signature. In our resting-state analysis, beta LSFP and RVT are not phase locked, suggesting an alternate mechanism at work. Nonetheless, beta power is thought to be the anti-alpha, being increased in states of cognitive concentration. Thus, the association between beta and RVT should be taken into account in resting-state EEG. Interestingly, in our study, we found that the parietal lobe is associated with the highest number of subjects exhibiting a negative lag (EEG leads RVT) (Fig. 6e), whereas the frontal lobe is associated with a larger number of subjects exhibiting positive lag (EEG lags RVT)(Fig. 6f). This observation is in line with the thought that the activity in the sensorimotor region (parietal) leads RVT, with the subsequent changes in cognition (frontal lobe) occurring later.

A novel finding amidst the inter-subject variability is that of preferential correlation polarities. While Yuan et al. reported predominantly positive correlations at negative lags (EEG leading RVT with a positive correlation) (Yuan et al., 2013), in Fig. 4 we noted that negative peak correlations are preferentially associated with negative lags. This is observed to different extents but nonetheless consistently across all EEG bands and lobes. However, as seen in Fig. 4a, the alpha band in the occipital lobe behaves rather differently than the other bands and in the other regions. That is, in the alpha region, EEG most often leads RVT with a positive correlation, consistent with the findings of Yuan et al.. The converse is also true, namely that positive lags seem to be preferentially associated with positive r values in these regions and bands. This finding is more observable in the parietal and occipital lobes than in the frontal lobe, though the differences amongst these lobes is not large. However, this observation is also more prominent in the theta band.

This finding of preferential polarity is very interesting and may help us interpret the significance of negative relative to positive correlations between EEG and RVT. Bearing in mind that negative lags indicate that EEG leads RVT, our data indicates that, in spite of high inter-subject variability, we are able to observe a pattern that is specific to the higher frequency EEG bands. As increases in EEG power in these bands are followed by a reduction in RVT in these brain regions, which in turn signify increased arousal, this observation is consistent with previous findings that indicate increased RVT during deep sleep (Pagliardini et al., 2013; Rostig et al., 2005). Moreover, widespread coupling between EEG and respiration is best observed in the theta band (Tort et al., 2018), where we also observe a stronger segregation between positive and negative EEG-RVT correlations by their lags. On the strength of this evidence, we postulate that positive and negative CCF peaks are both valid, and can be interpreted in different ways. On one hand, as described in earlier sections, respiration changes are known to lead to changes in higher-order function that involves the alpha wave (Heck et al., 2016). On the other hand, what we are observing is inevitably a part of a cycle in which respiration feeds back into neuronal regulation, which in turn further regulates respiration. The entrainment of RVT by EEG and vice versa merit further study.

The cross-spectral coherence analyses revealed that the frequencies of peak coherence between EEG alpha/delta LSFP and RVT are largely found at ≤0.02 Hz, whereas the theta-RVT and beta-RVT associations appear to peak at ≥0.02 Hz. Both ranges are frequencies of interest for rs-fMRI analyses.

### Resting-state respiration and neural activity: Temporal and intersubject variability

Unsurprisingly, the associations between RVT and EEG LSFP are non-stationary, as illustrated by the wavelet-cross spectrum (WCS) plots (Fig. 9). Despite the rhythmic nature of EEG signals and the largely cyclic nature of RVT, the variability in their relationship is quite substantial. These WCS plots provide a sense of the temporal variability in terms of the strength and direction of EEG-RVT coupling, and suggest that measurements of the coupling taken at different times or in different sessions will not necessarily be similar.

As demonstrated by Yuan et al. (Yuan et al., 2013), one of the key factors modulating the inter-subject variability in the RVT-EEG relationship is likely to be arousal. To test this possibility, we computed a vigilance measure based on the ratio of the fast and slow rhythms (Liu and Falahpour, 2020). However, we showed that RVT-EEG associations are largely independent of vigilance.

### Implications for resting-state fMRI

The frequency of peak coherence between RVT and EEG in different frequency bands are all directly within the key frequency range for rs-fMRI, even after taking into account the hemodynamic lag that relates EEG to fMRI (de Munck et al., 2007). Of course, within this range, our results could be influenced by artifacts, including movement and MRI artifacts. However, as a result of our diligent denoising and manual quality-control procedure, we have confidence that these results are not primarily driven by artifacts.

Here we demonstrate that the EEG-RVT connection is generalizable across multiple frequency bands and brain regions. However, we have identified at least two types of interactions. In the case of EEG leading RVT, the mechanism is perhaps better understood, particularly as the sensorimotor, supplementary motor areas as well as the brainstem and thalamus are a part of respiratory control. In the context of rs-fMRI, the sensorimotor network is sometimes studied, but much more often, networks related to higher brain function, including the default-mode network, the frontal-attention network and the salience network, are the objective of investigation. For the purpose of studying higher brain function, it is likely that the EEG activity that is induced by RVT is of greater interest. After all, the feedback between respiration and cortical activity through ANS regulation is the mechanism that has direct bearing on cognition and emotional regulation. Thus, by removing the influence of RVT on the rs-fMRI signal, we may risk losing relevant effects related to higher brain function.

### Limitations and future research

There are a number of limitations in this study, chief among them being the simplification of the RVT-EEG relationship. Although we probed the nonstationarity of this relationship, we found it more difficult to interpret than a stationary approximation (as represented by the cross spectral peaks). Moreover, in this initial proof-of-concept study, we did not consider non-linearities in the EEG-RVT association (Morelli et al., 2018), which are likely beyond the scope of this study but will be pursued in our future work.

Furthermore, we opted to exclude the gamma band, due to signal-quality reasons, as mentioned in Methods. Nonetheless, the gamma band is of particular interest to resting-state fMRI. Thus, despite the well-known artifactual challenges in investigating the gamma band (Muthukumaraswamy, 2013), methodological refinements to that end will also be pursued in our future work.

## Acknowledgments

This research is funded by the Canadian Institutes of Health Research (CIHR) and the Sandra Rotman Foundation.

## Supplementary Materials

**Table A1.** Results of stationarity tests for RVT and EEG global field power (GFP) for different subjects.

| Subject ID | for RVT |  | EEG_GFP |  |
| --- | --- | --- | --- | --- |
|  | P-value | Decision | P-value | Decision |
| S1 | 0.1 | stationary | 0.01 | Not stationary |
| S2 | 0.01 | nonstationary | 0.1 | stationary |
| S3 | 0.01 | nonstationary | 0.1 | stationary |
| S4 | 0.1 | stationary | 0.1 | stationary |
| S5 | 0.07 | stationary | 0.1 | stationary |
| S6 | 0.01 | nonstationary | 0.02 | nonstationary |
| S7 | 0.01 | nonstationary | 0.1 | stationary |
| S8 | 0.1 | stationary | 0.01 | nonstationary |
| S9 | 0.03 | nonstationary | 0.1 | stationary |
| S10 | 0.1 | stationary | 0.02 | nonstationary |
| S11 | 0.02 | nonstationary | 0.01 | nonstationary |
| S12 | 0.04 | nonstationary | 0.01 | nonstationary |
| S13 | 0.04 | nonstationary | 0.03 | nonstationary |
| S14 | 0.01 | nonstationary | 0.02 | nonstationary |
| S15 | 0.01 | nonstationary | 0.01 | nonstationary |
| S16 | 0.1 | stationary | 0.01 | nonstationary |
| S17 | 0.01 | nonstationary | 0.01 | nonstationary |

**Table A2.**
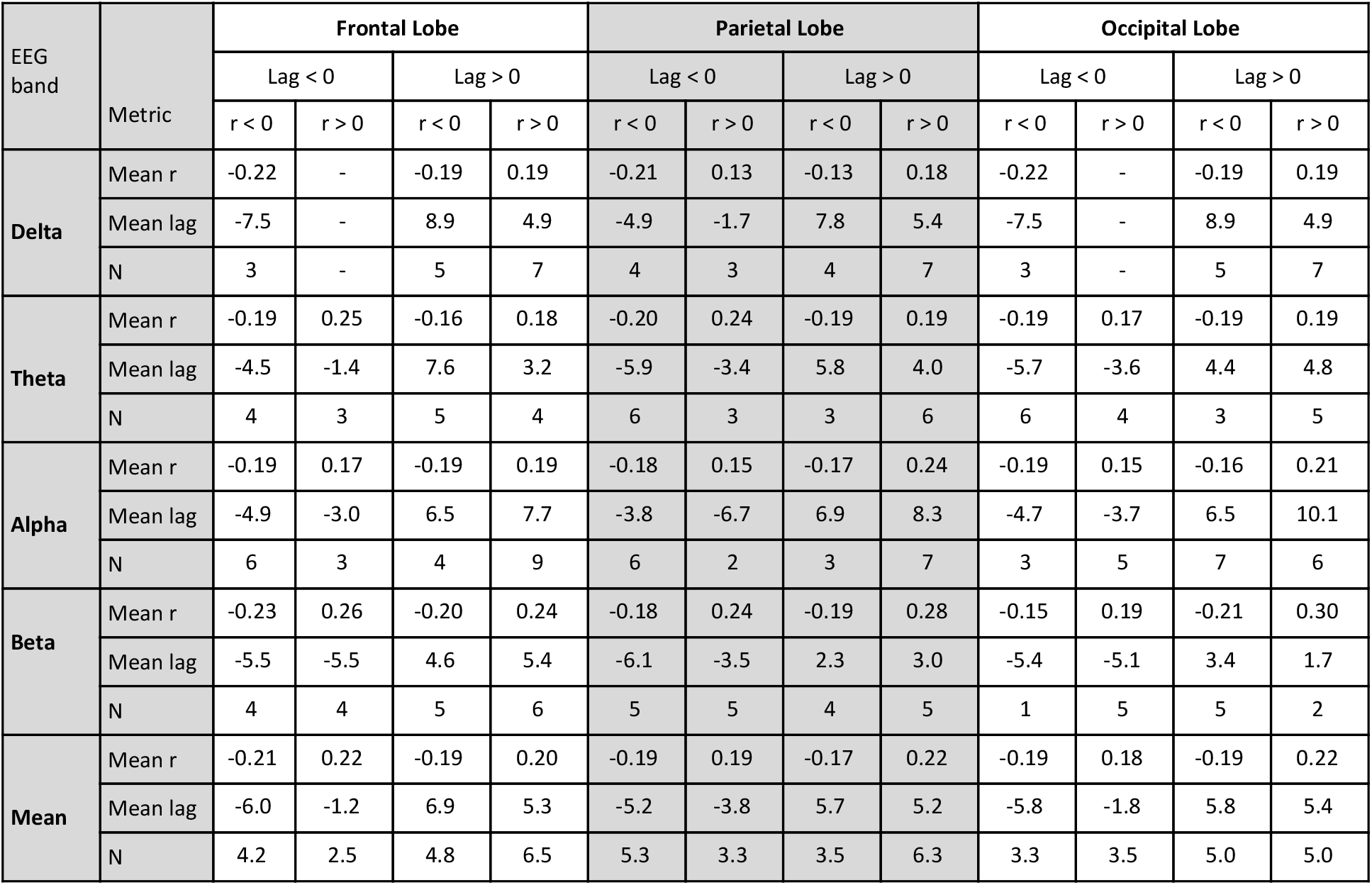
Summary of lag and correlations by sign and the lag.

| EEG band | Metric | Frontal Lobe |  |  |  | Parietal Lobe |  |  |  | Occipital Lobe |  |  |  |
| --- | --- | --- | --- | --- | --- | --- | --- | --- | --- | --- | --- | --- | --- |
|  |  | Lag < 0 |  | Lag > 0 |  | Lag < 0 |  | Lag > 0 |  | Lag < 0 |  | Lag > 0 |  |
|  |  | r < 0 | r > 0 | r < 0 | r > 0 | r < 0 | r > 0 | r < 0 | r > 0 | r < 0 | r > 0 | r < 0 | r > 0 |
| Delta | Mean r | -0.22 | - | -0.19 | 0.19 | -0.21 | 0.13 | -0.13 | 0.18 | -0.22 | - | -0.19 | 0.19 |
|  | Mean lag | -7.5 | - | 8.9 | 4.9 | -4.9 | -1.7 | 7.8 | 5.4 | -7.5 | - | 8.9 | 4.9 |
|  | N | 3 | - | 5 | 7 | 4 | 3 | 4 | 7 | 3 | - | 5 | 7 |
| Theta | Mean r | -0.19 | 0.25 | -0.16 | 0.18 | -0.20 | 0.24 | -0.19 | 0.19 | -0.19 | 0.17 | -0.19 | 0.19 |
|  | Mean lag | -4.5 | -1.4 | 7.6 | 3.2 | -5.9 | -3.4 | 5.8 | 4.0 | -5.7 | -3.6 | 4.4 | 4.8 |
|  | N | 4 | 3 | 5 | 4 | 6 | 3 | 3 | 6 | 6 | 4 | 3 | 5 |
| Alpha | Mean r | -0.19 | 0.17 | -0.19 | 0.19 | -0.18 | 0.15 | -0.17 | 0.24 | -0.19 | 0.15 | -0.16 | 0.21 |
|  | Mean lag | -4.9 | -3.0 | 6.5 | 7.7 | -3.8 | -6.7 | 6.9 | 8.3 | -4.7 | -3.7 | 6.5 | 10.1 |
|  | N | 6 | 3 | 4 | 9 | 6 | 2 | 3 | 7 | 3 | 5 | 7 | 6 |
| Beta | Mean r | -0.23 | 0.26 | -0.20 | 0.24 | -0.18 | 0.24 | -0.19 | 0.28 | -0.15 | 0.19 | -0.21 | 0.30 |
|  | Mean lag | -5.5 | -5.5 | 4.6 | 5.4 | -6.1 | -3.5 | 2.3 | 3.0 | -5.4 | -5.1 | 3.4 | 1.7 |
|  | N | 4 | 4 | 5 | 6 | 5 | 5 | 4 | 5 | 1 | 5 | 5 | 2 |
| Mean | Mean r | -0.21 | 0.22 | -0.19 | 0.20 | -0.19 | 0.19 | -0.17 | 0.22 | -0.19 | 0.18 | -0.19 | 0.22 |
|  | Mean lag | -6.0 | -1.2 | 6.9 | 5.3 | -5.2 | -3.8 | 5.7 | 5.2 | -5.8 | -1.8 | 5.8 | 5.4 |
|  | N | 4.2 | 2.5 | 4.8 | 6.5 | 5.3 | 3.3 | 3.5 | 6.3 | 3.3 | 3.5 | 5.0 | 5.0 |

